## Supplementary figures for "Spinal premotor interneurons controlling antagonistic muscles are spatially intermingled"

### Figure 3 figure supplement 1

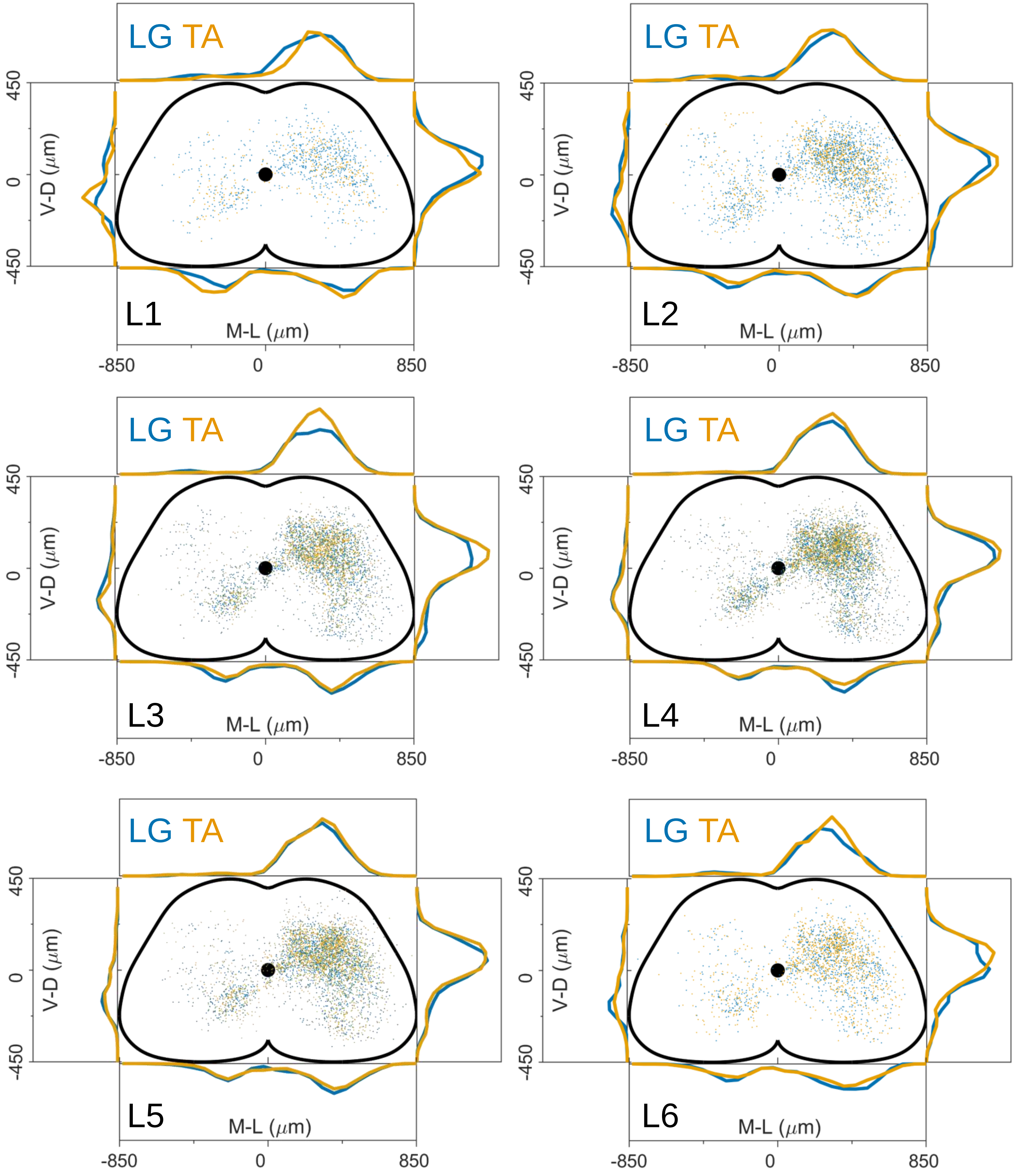

Figure 3 figure supplement 2

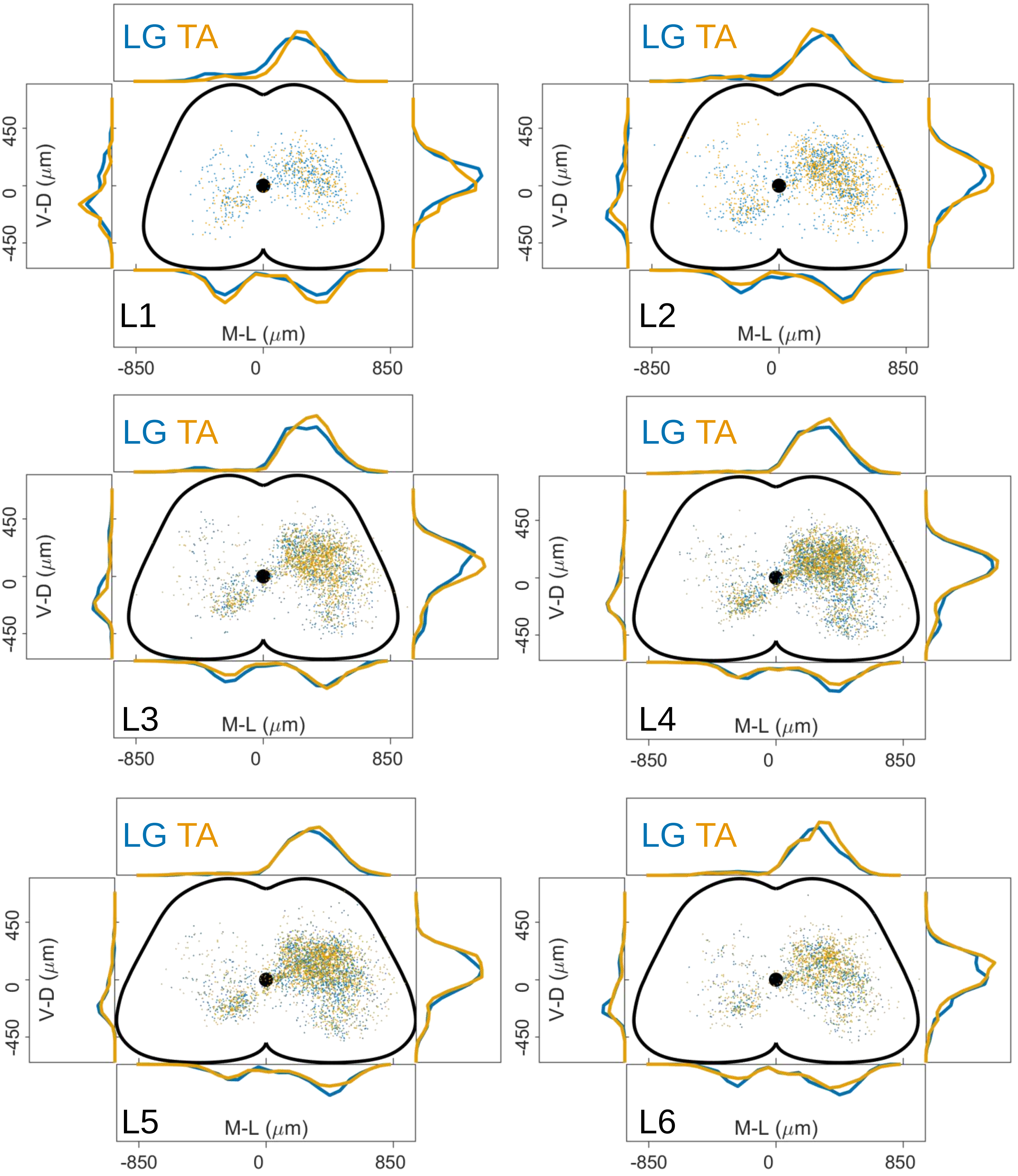

### Figure 3 figure supplement 3

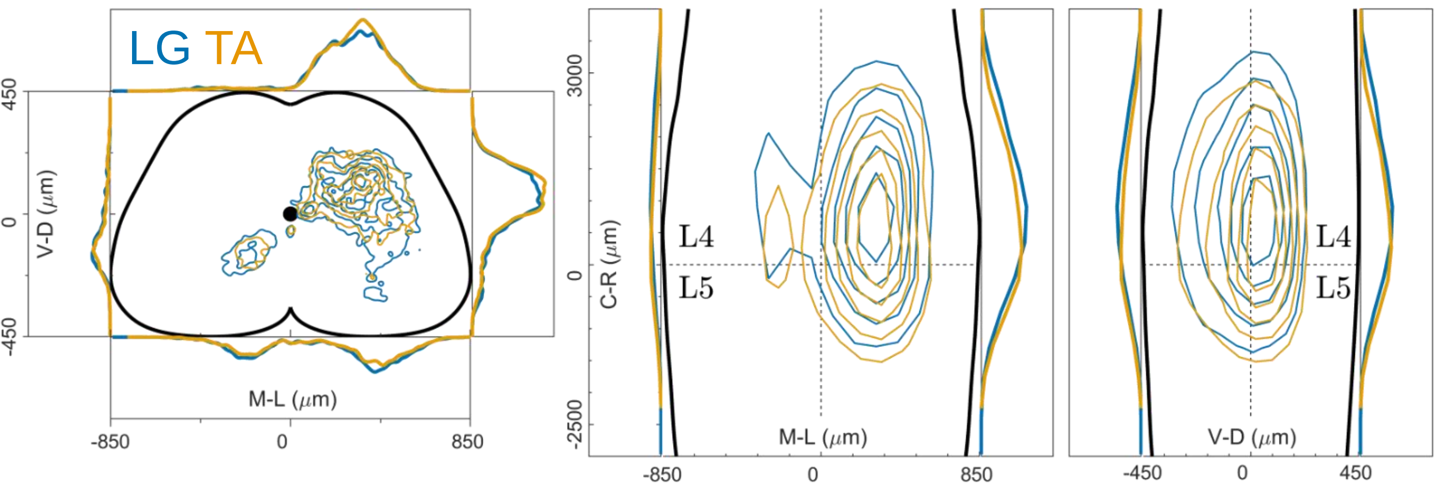

### Figure 4 figure supplement 1

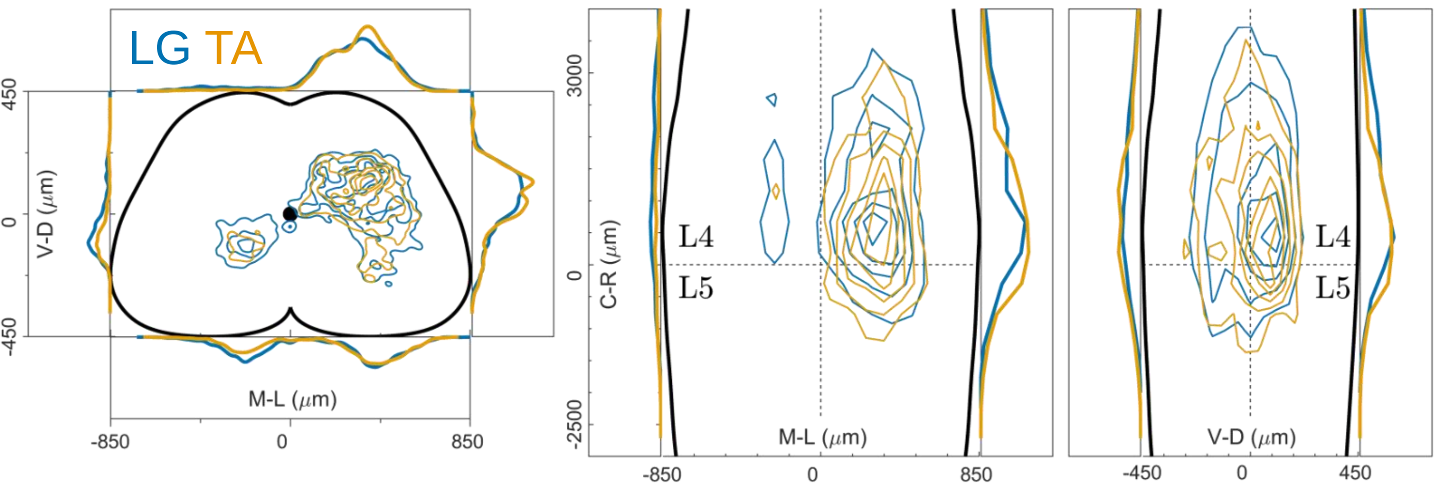

Figure 4 Figure supplement 2

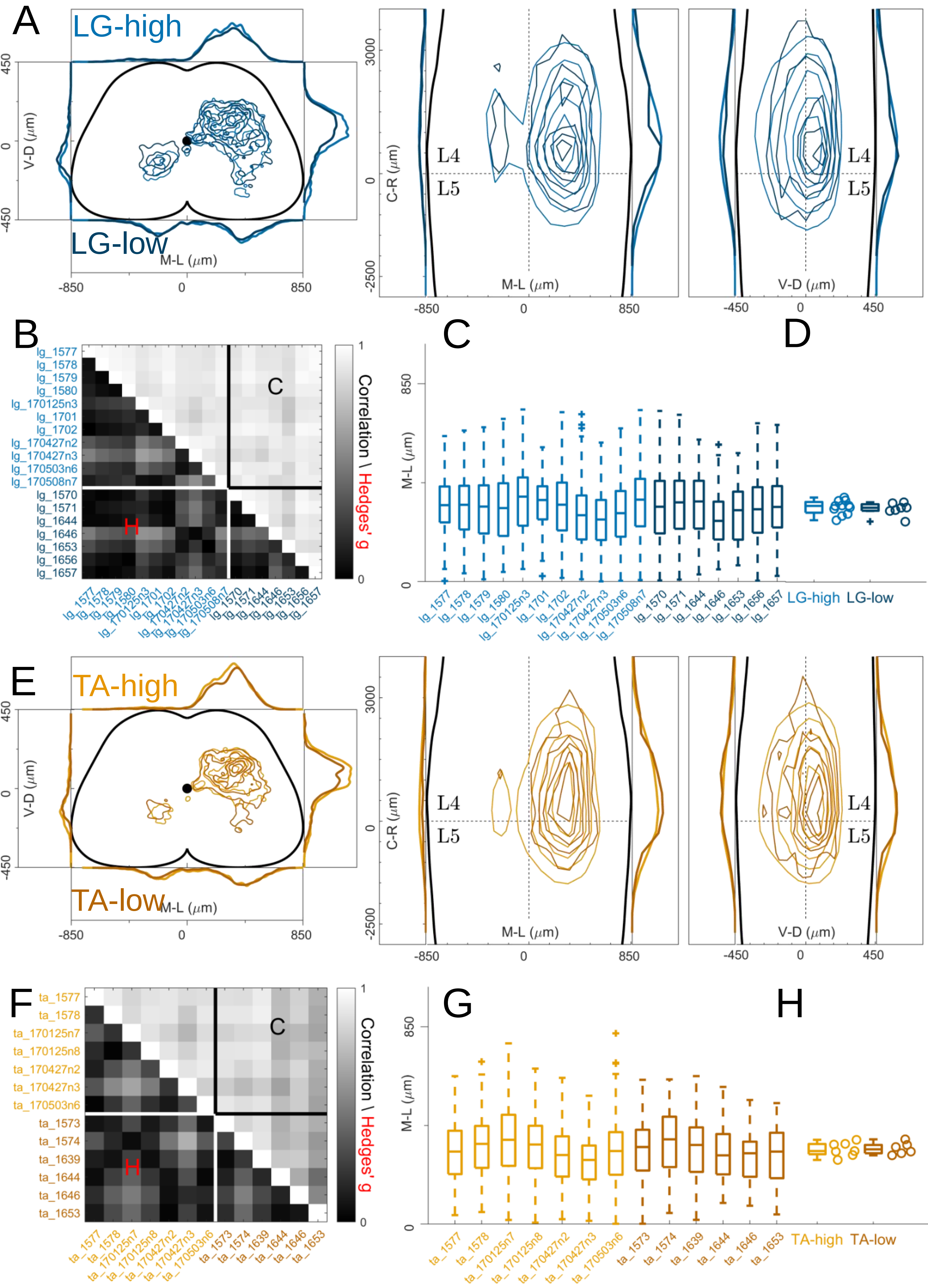

Figure 4 Figure supplement 3

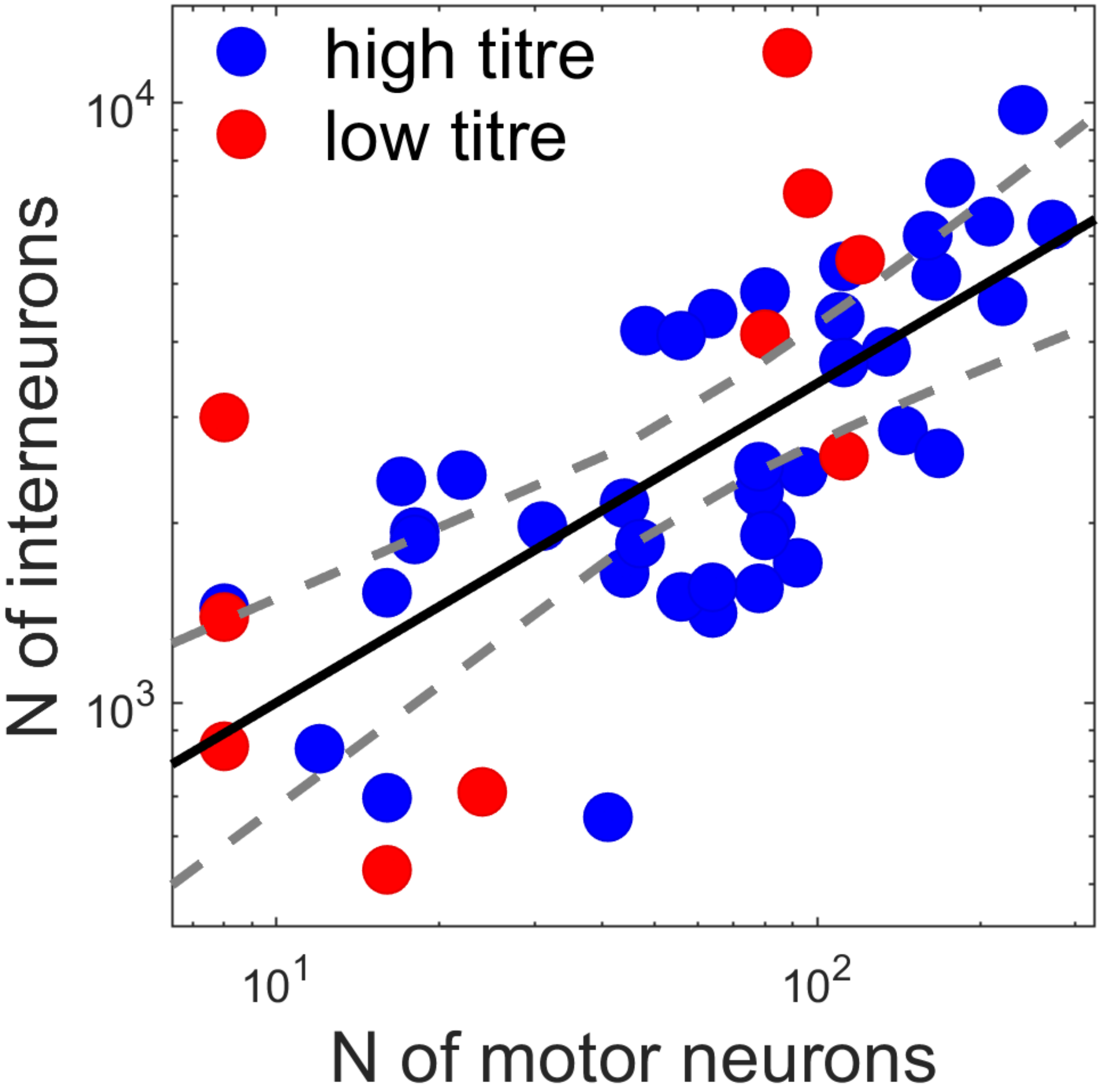

### Figure 7 Figure supplement 1

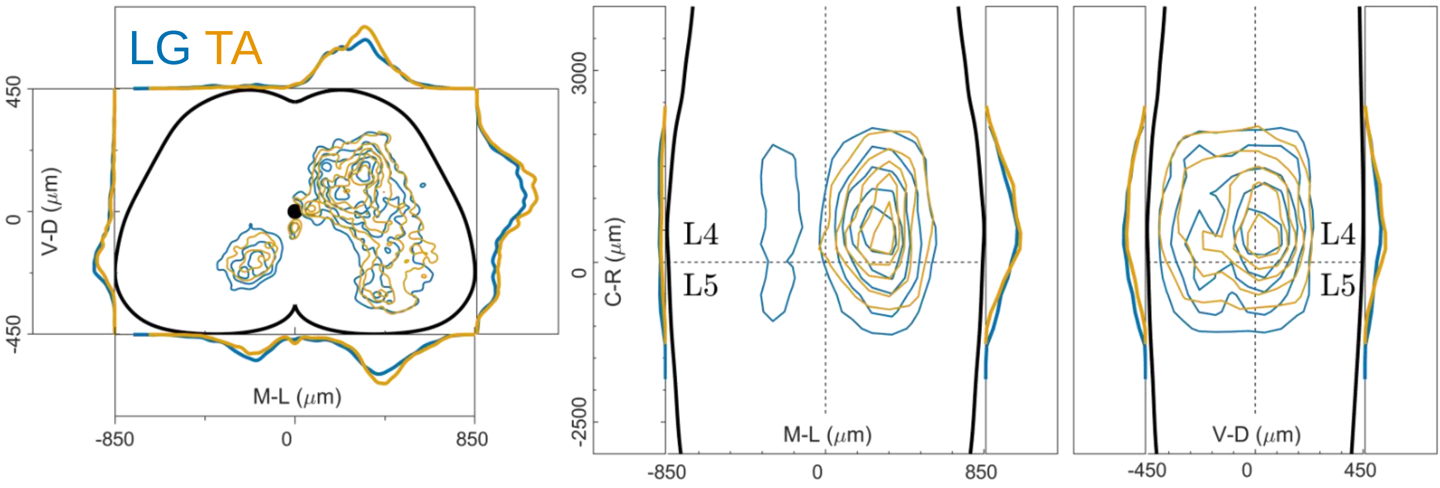

### Figure 12 figure supplement 1

Double injections UCL

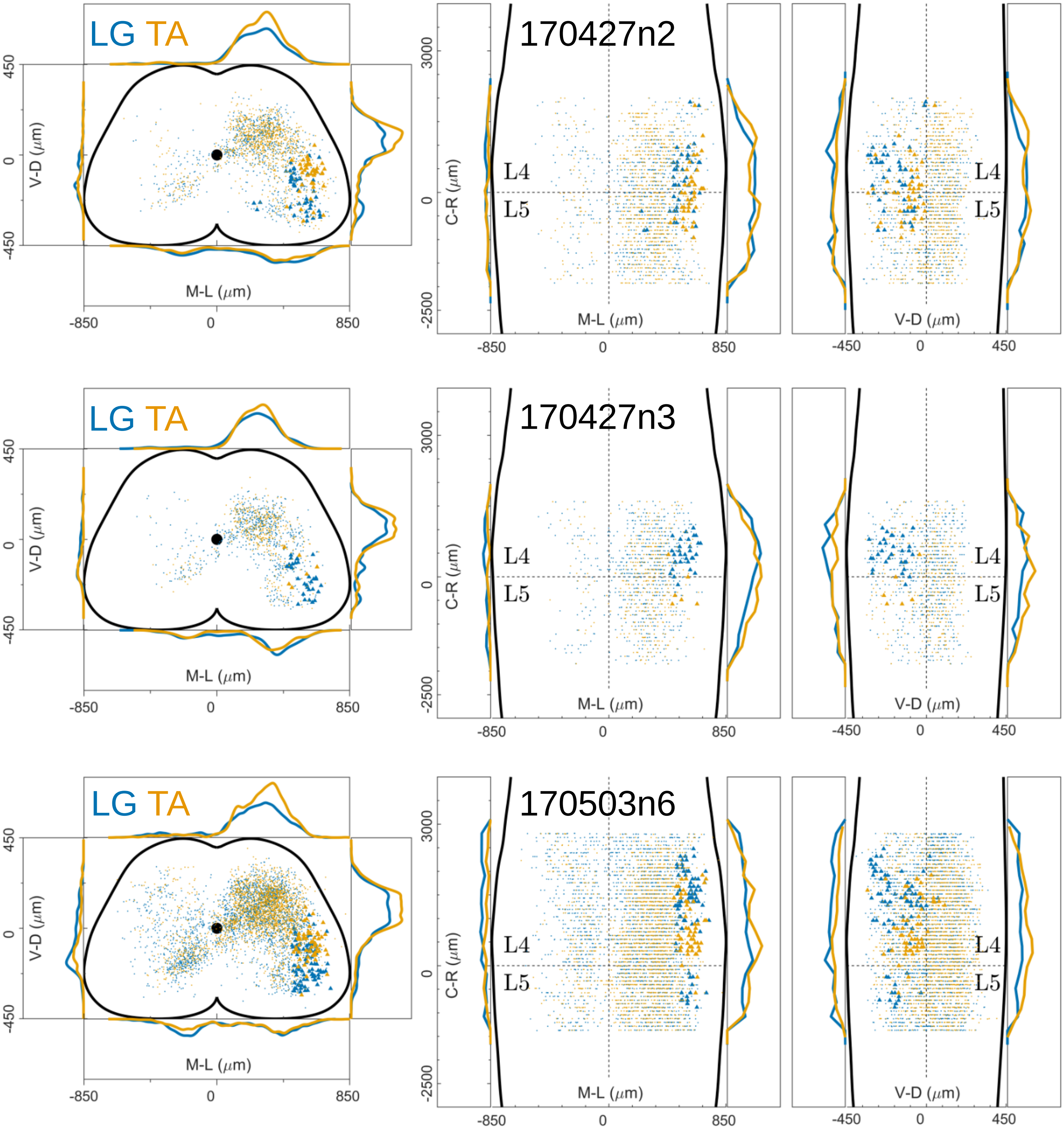

### Figure 12 figure supplement 2

double injections UoG

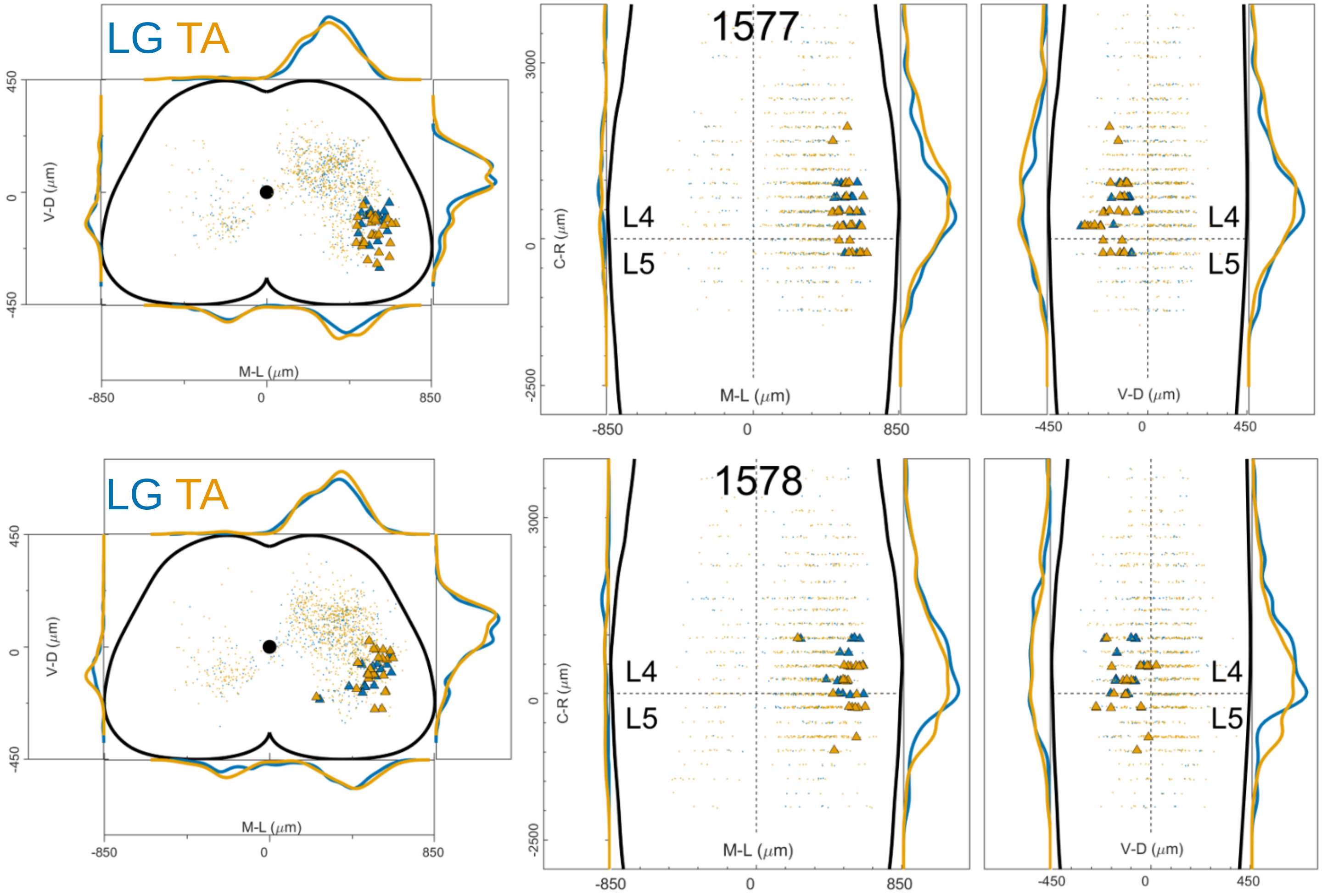

### Figure 12 figure supplement 3

Double injections UCL

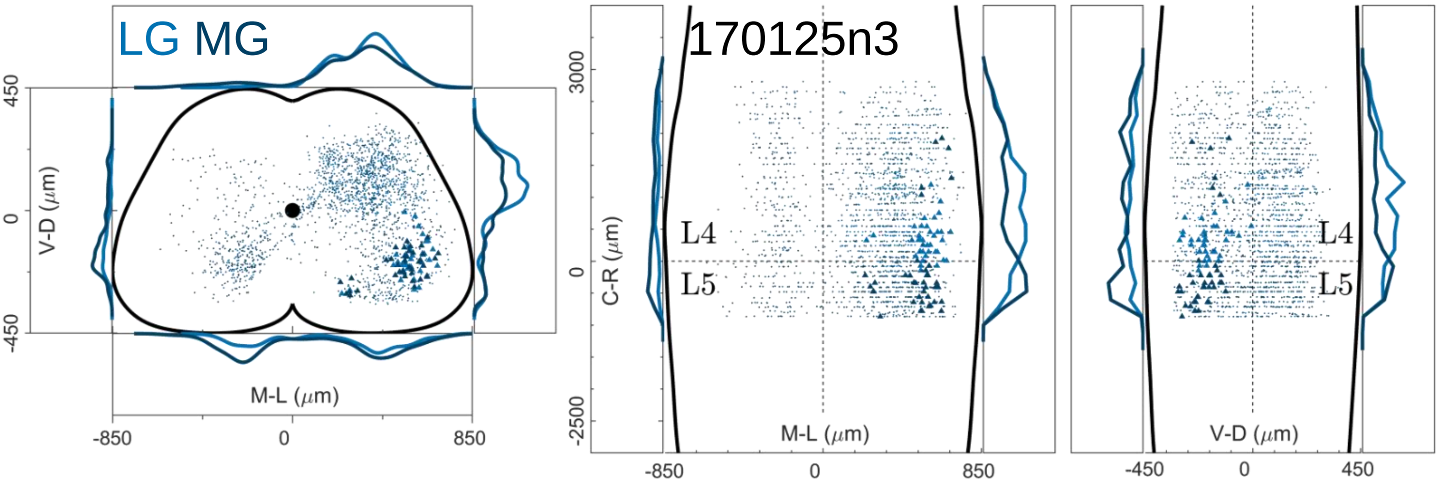


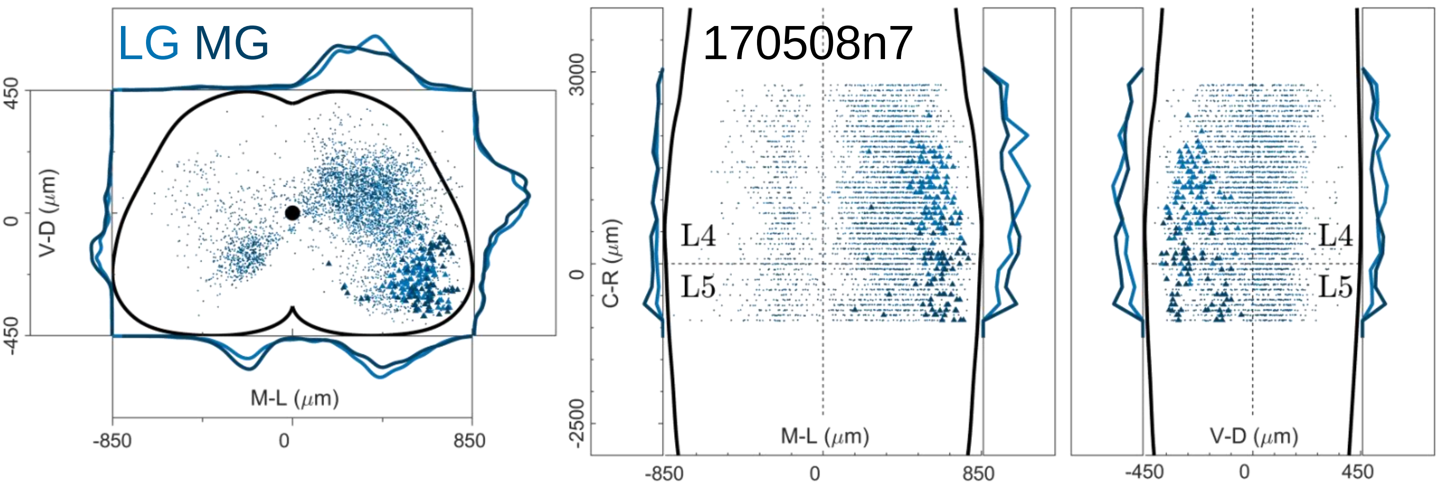

### Figure 12 figure supplement 4

Double injections UoG

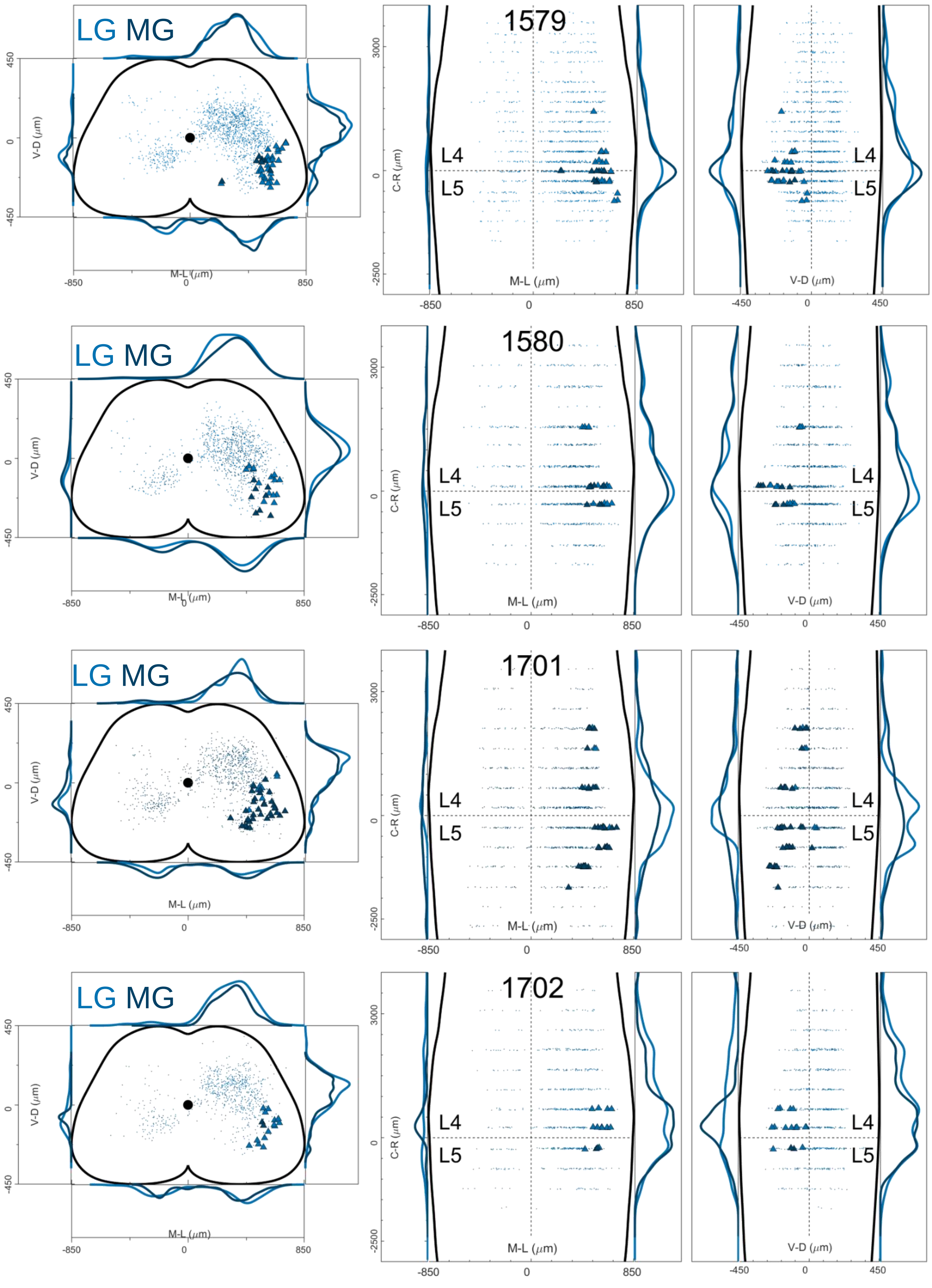

Figure 12 figure supplement 5

Double injections UCL

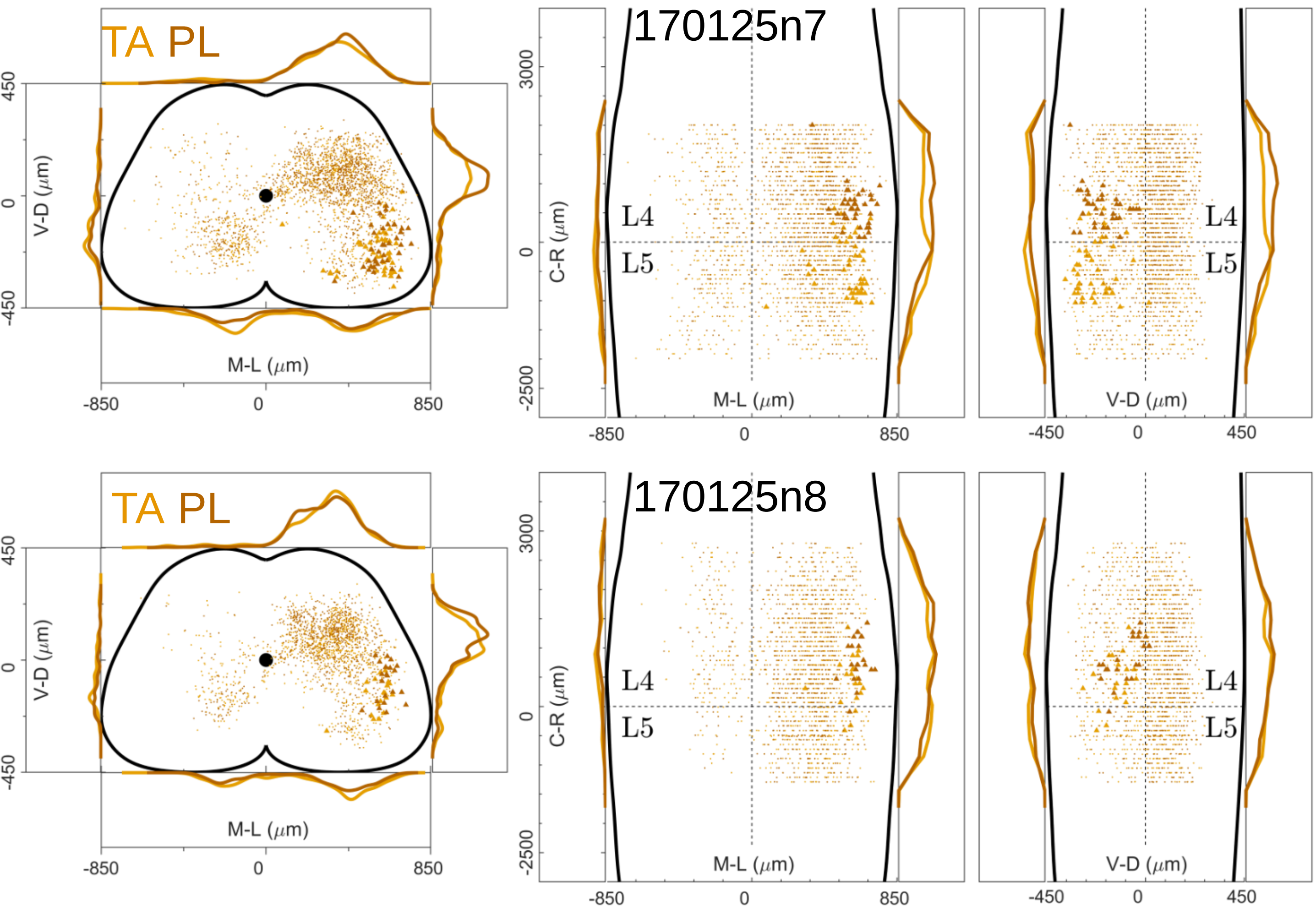

Figure 12 figure supplement 6 Single injections UoG

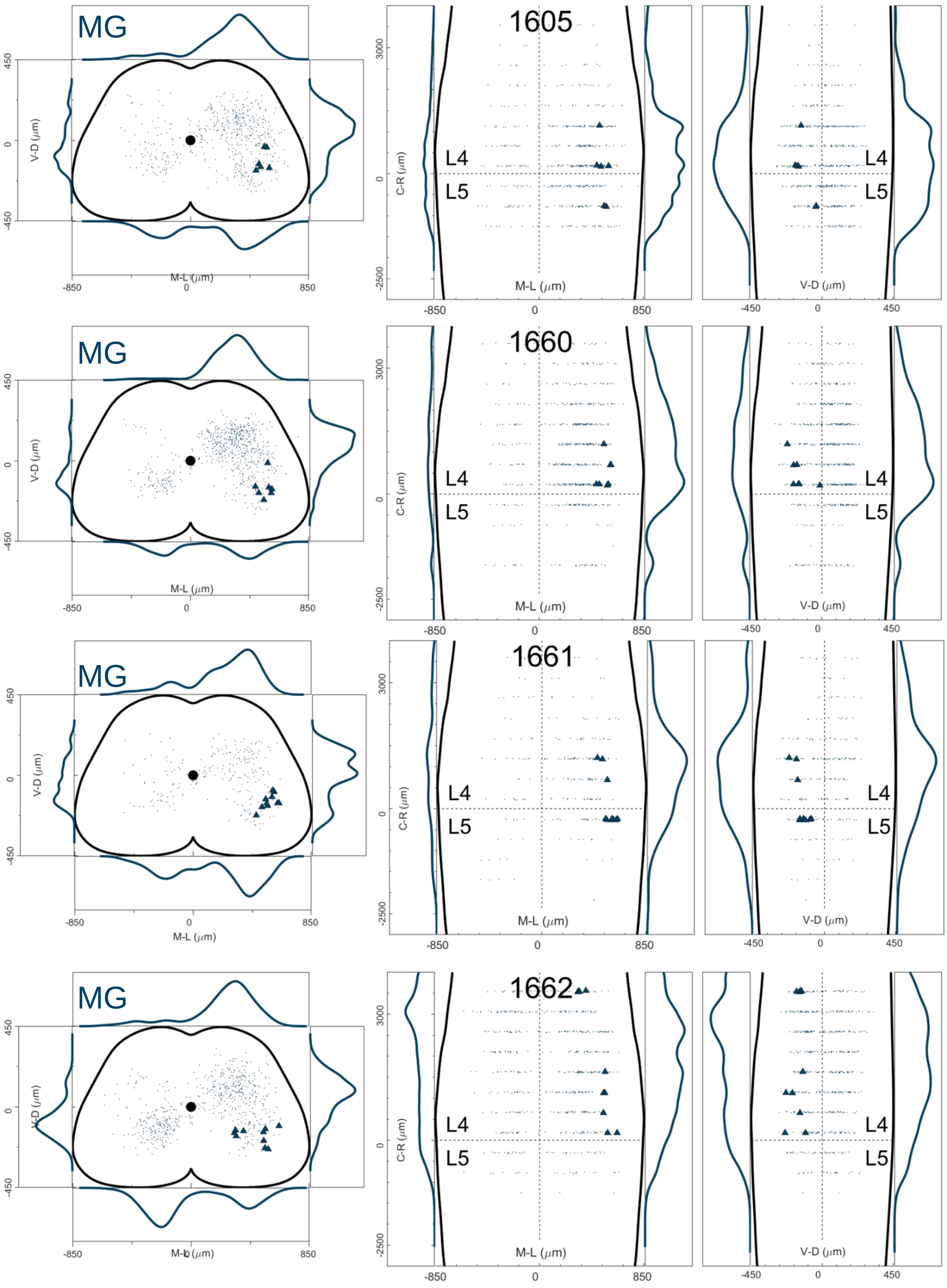

Figure 12 figure supplement 7

Single injections UoG

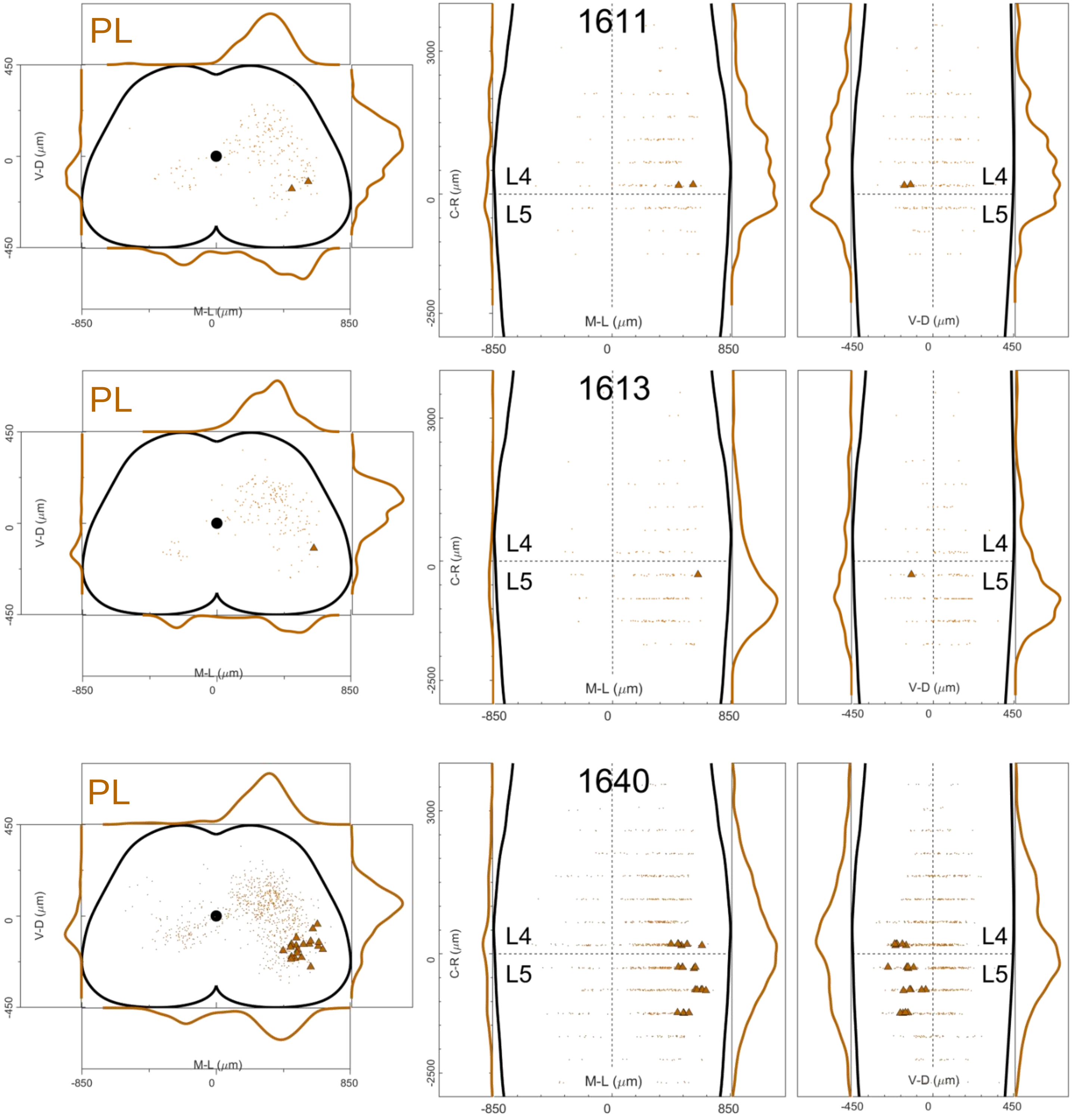

Figure 12 figure supplement 8

Low titre double injections UoG

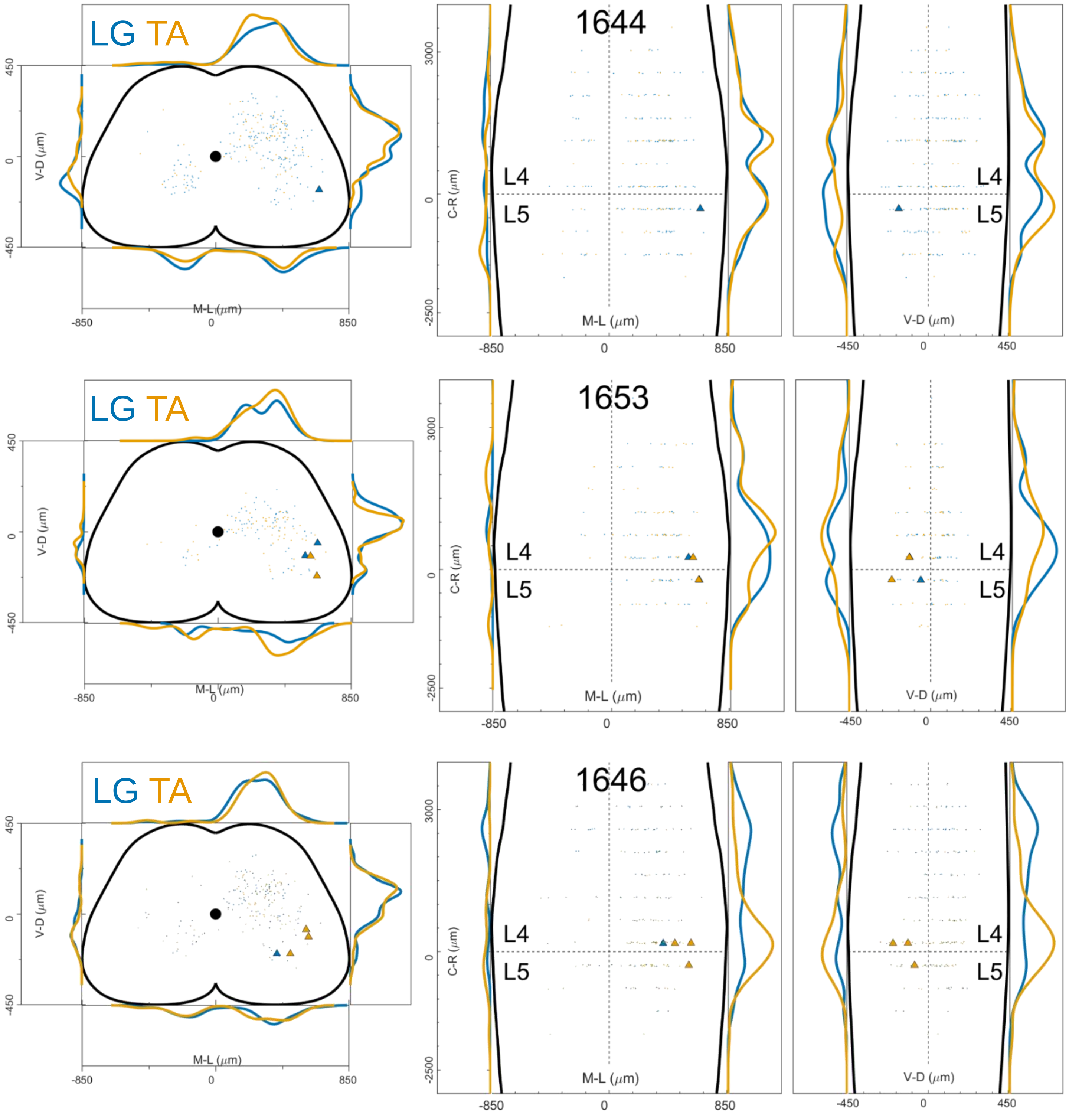

Figure 12 figure supplement 9 Low titre single injections UoG

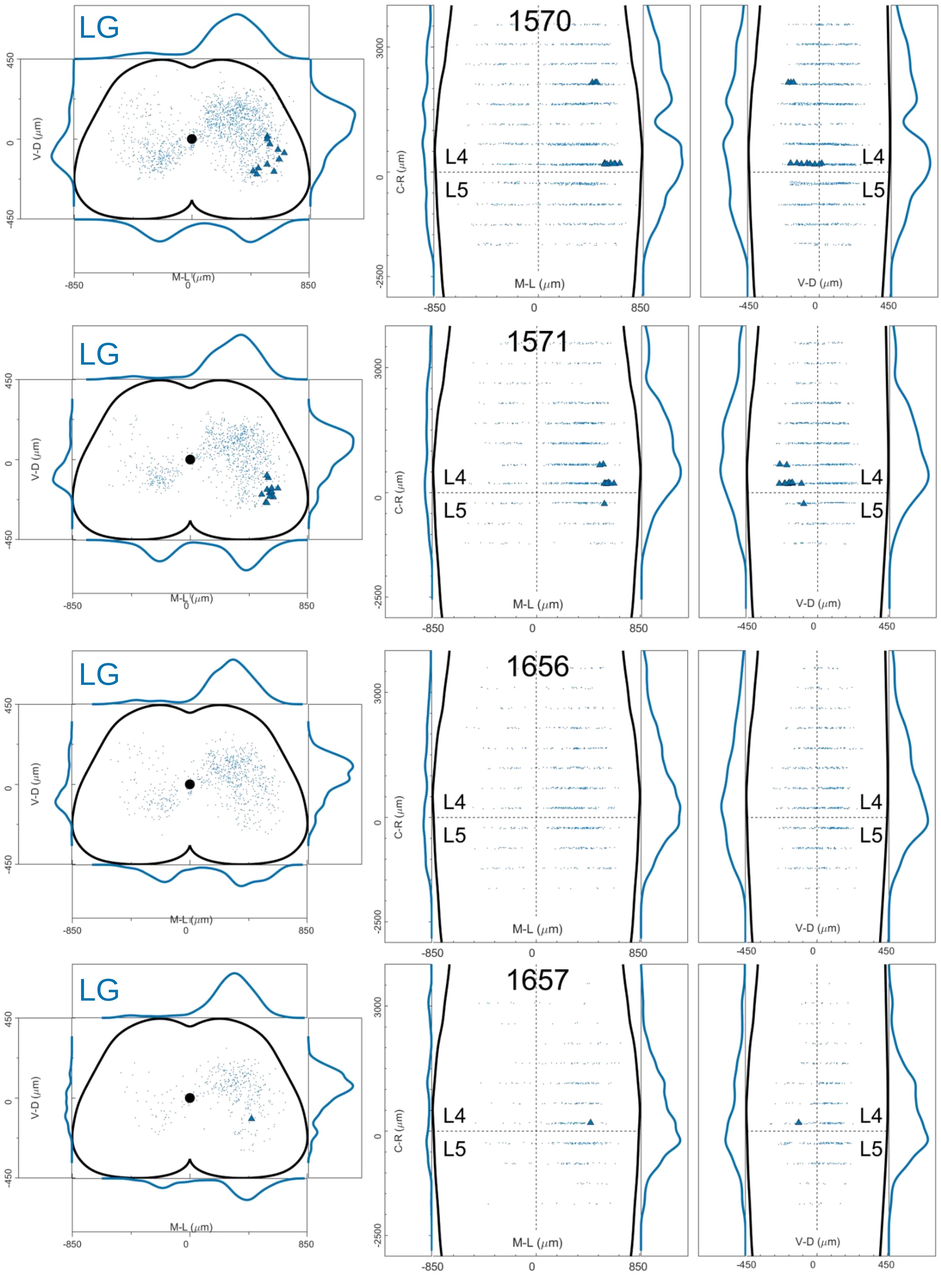

Figure 12 figure supplement 10 Low titre single injections UoG

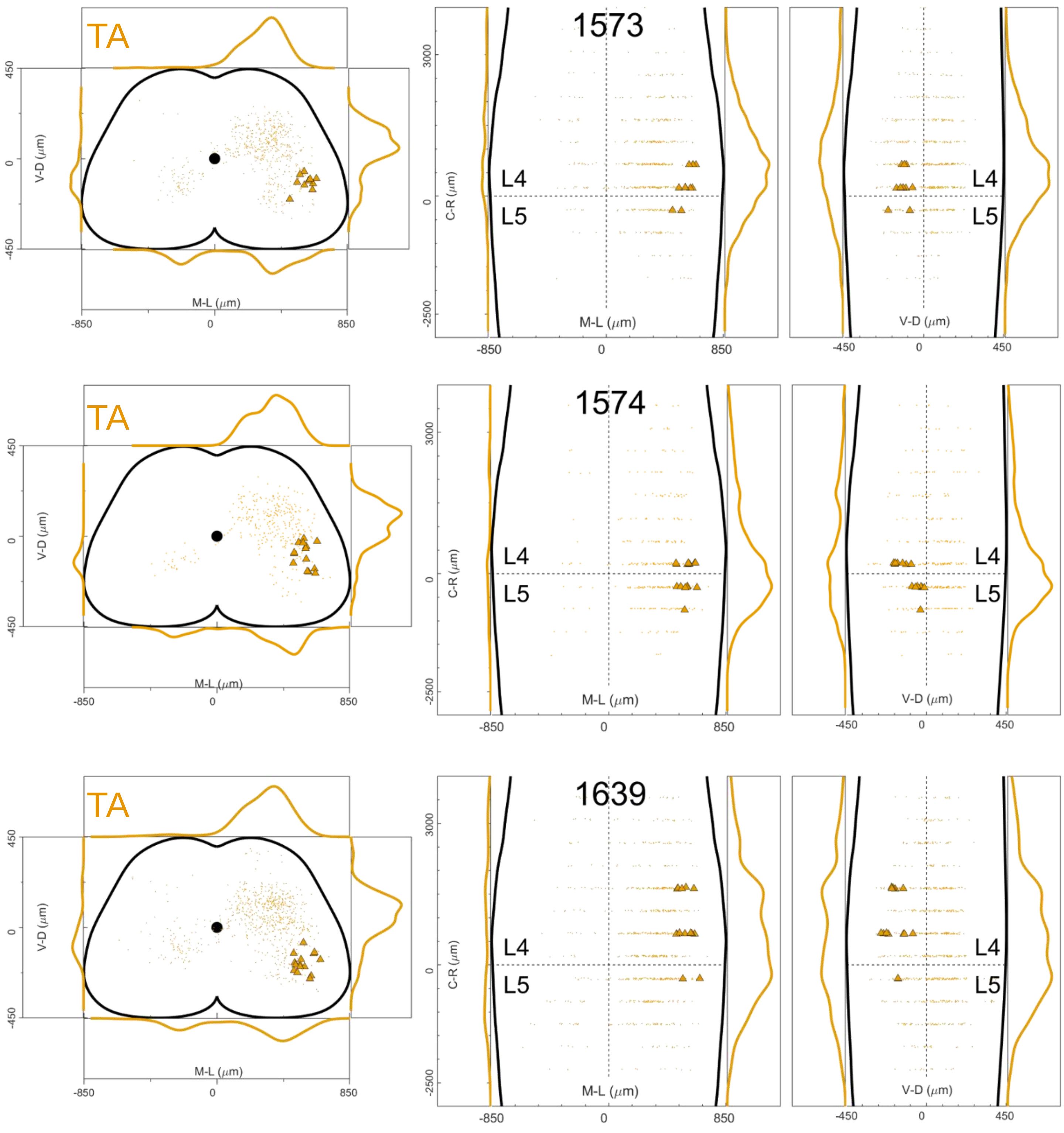

Figure 12 figure supplement 11 Single injections MDC

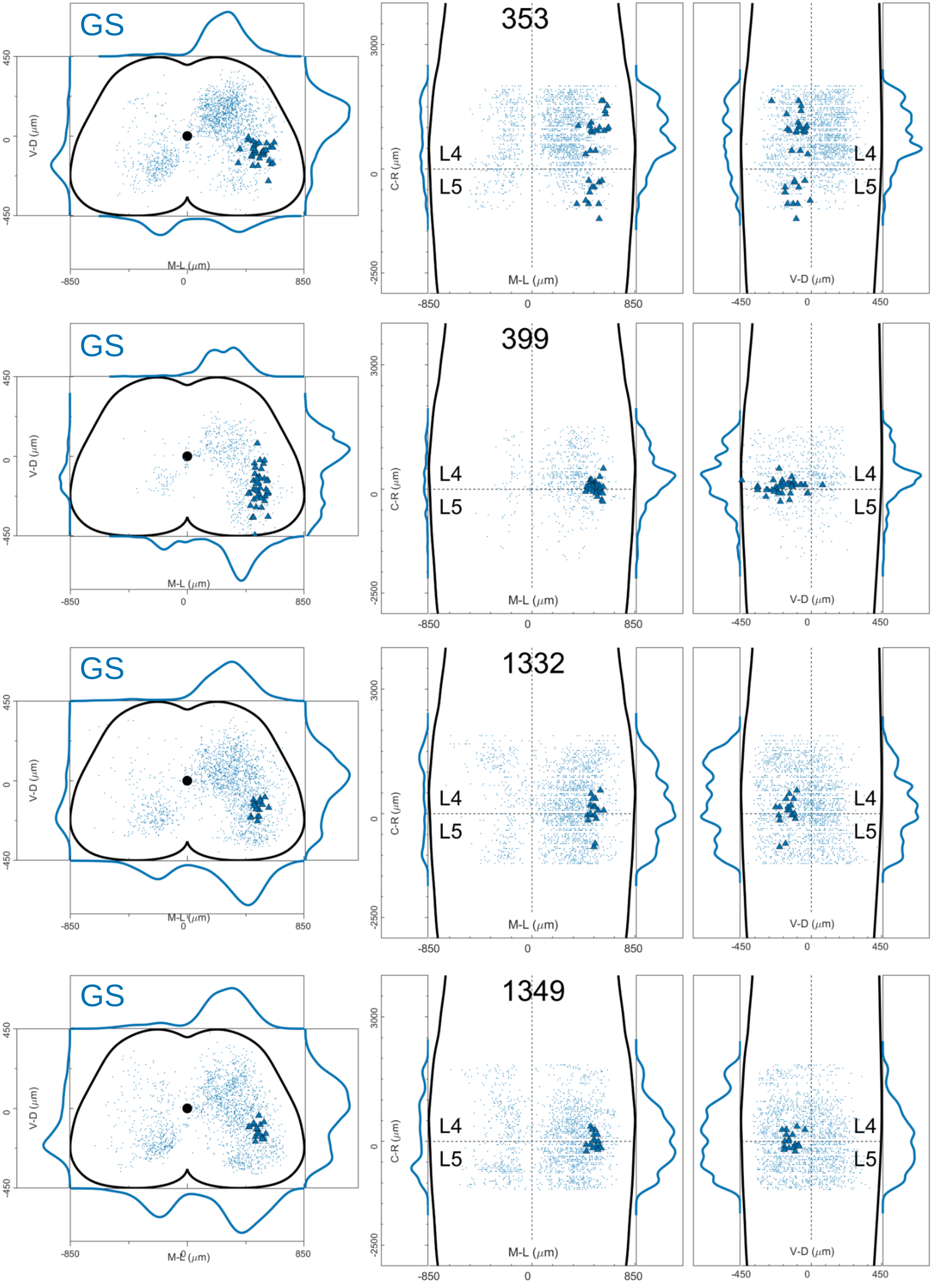

### Figure 12 figure supplement 12

Single injections MDC

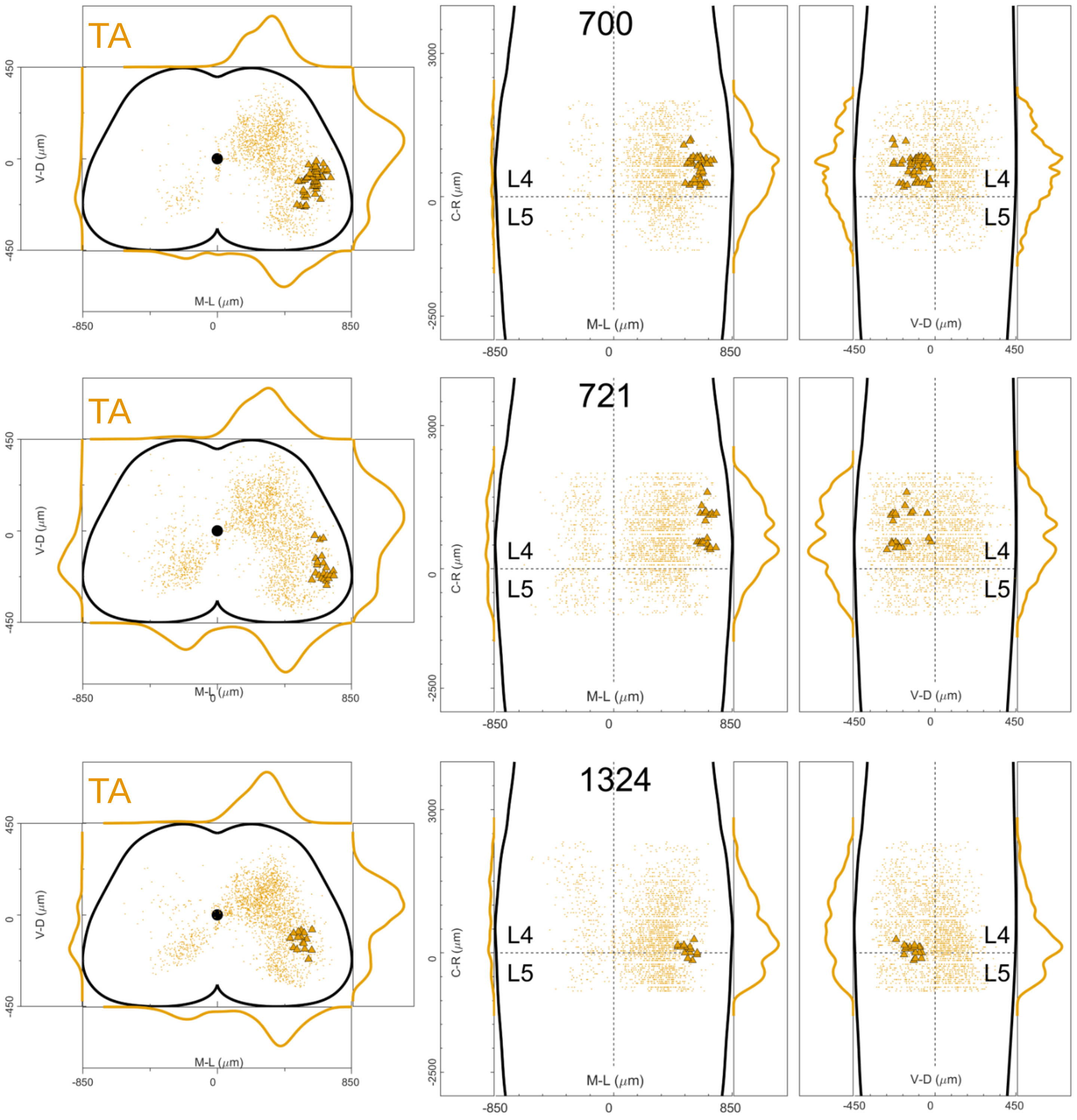

### Figure 12 figure supplement 13

injections AAV-G_wt Salk

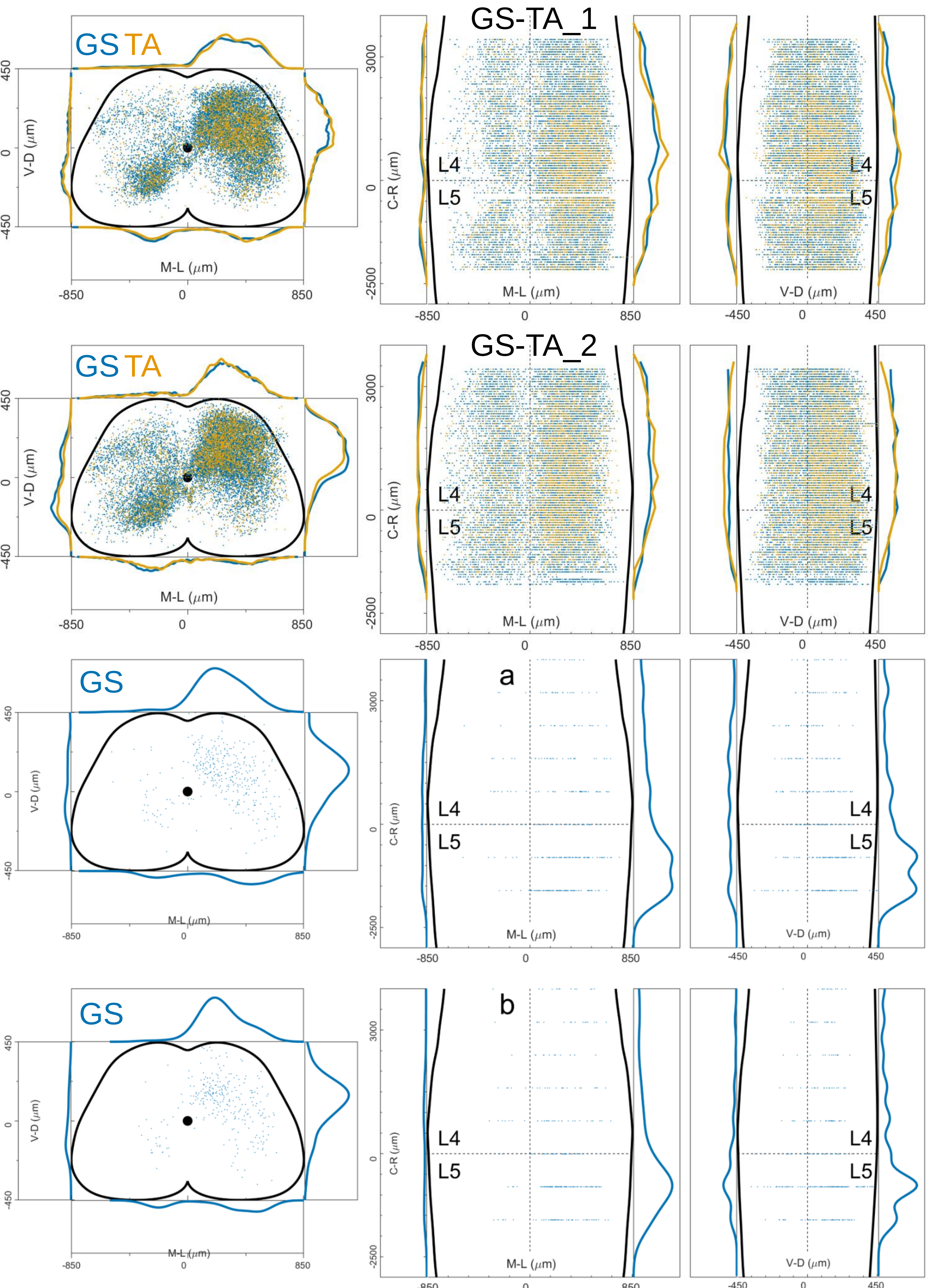

Figure 12 figure supplement 14 Injections AAV-G-Flex_ChAT Salk

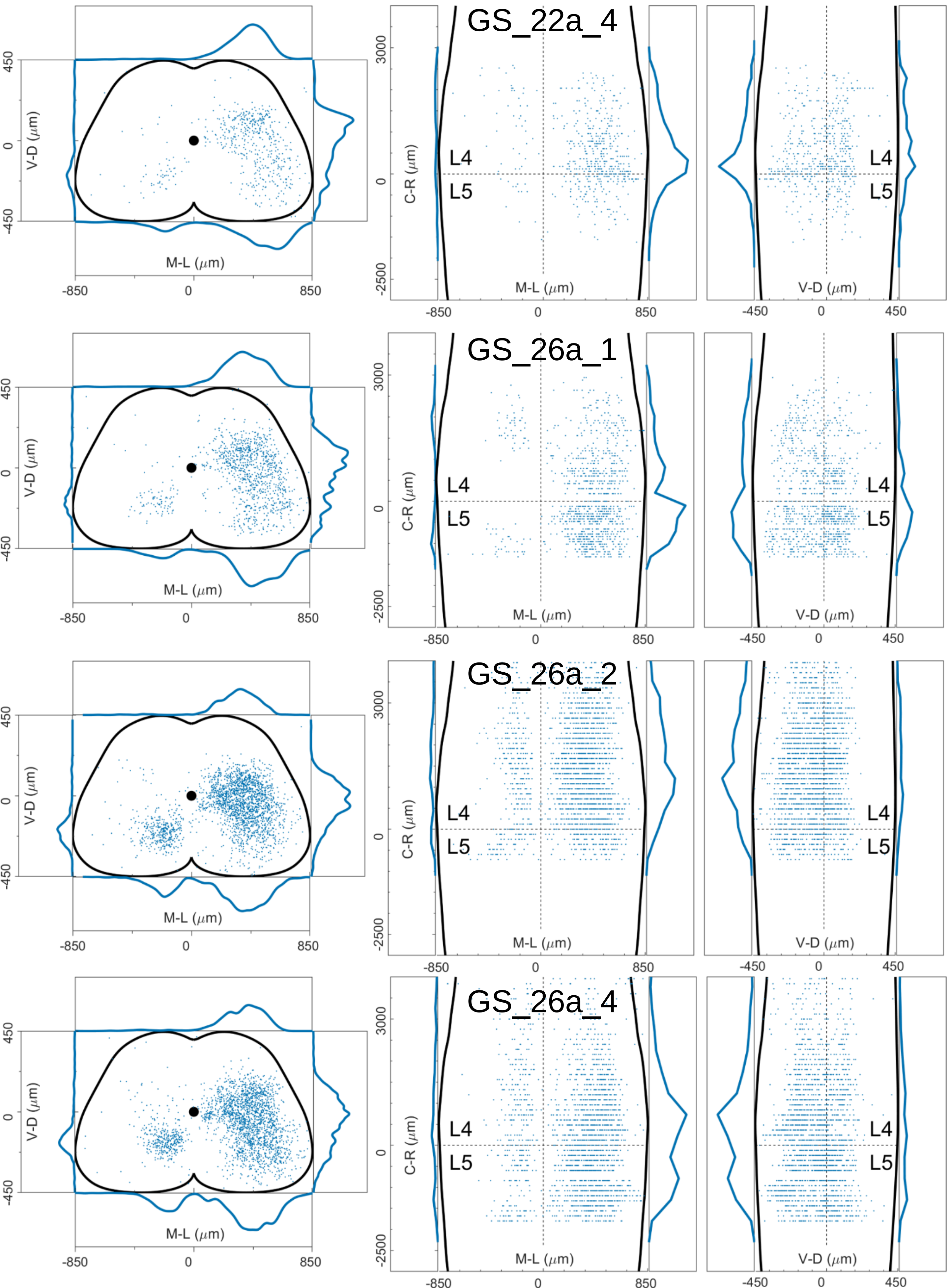

Figure 12 figure supplement 15 Injections AAV-G-Flex_ChAT Salk

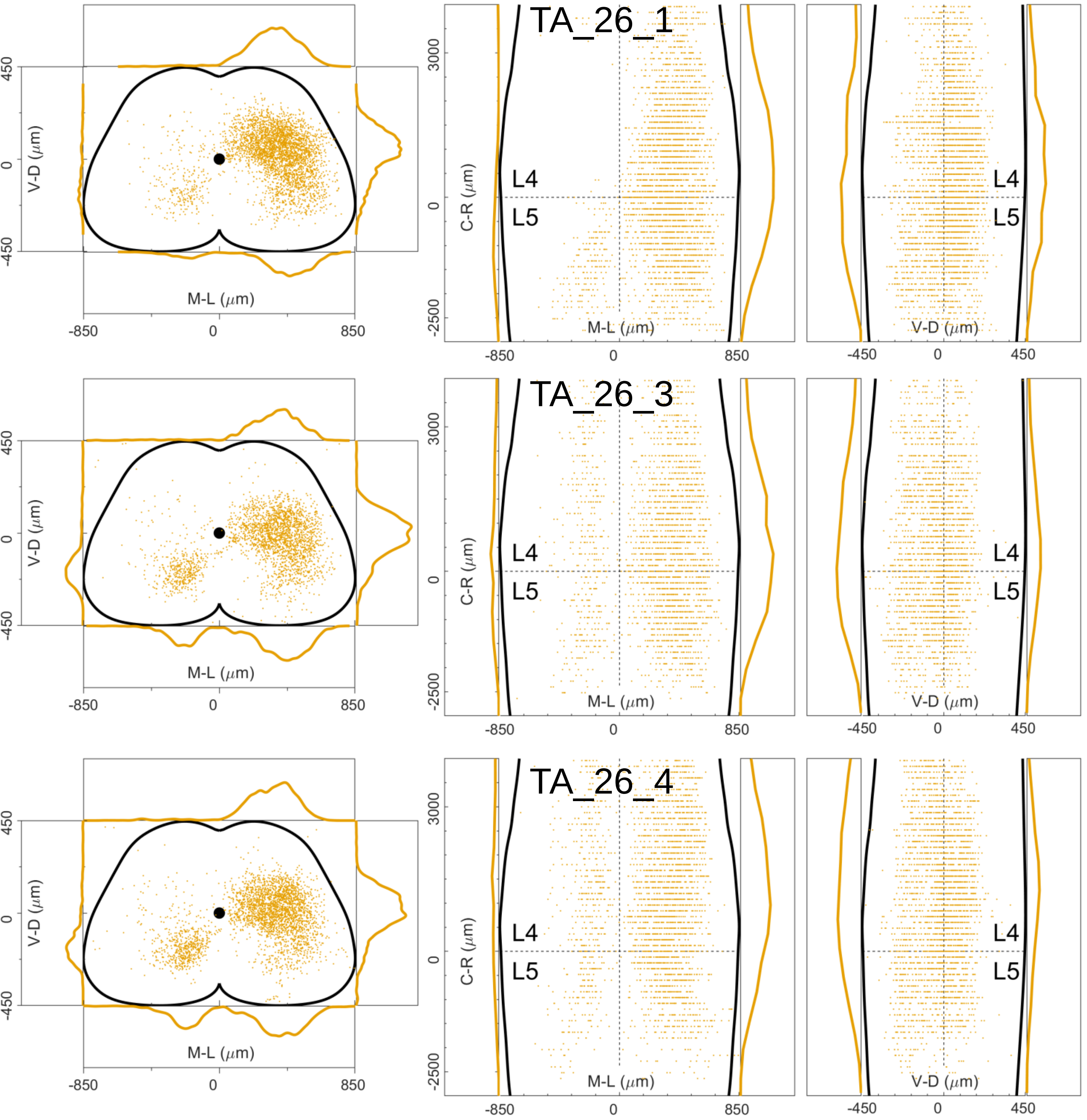

Figure 12 figure supplement 16

Double injections PRV Salk

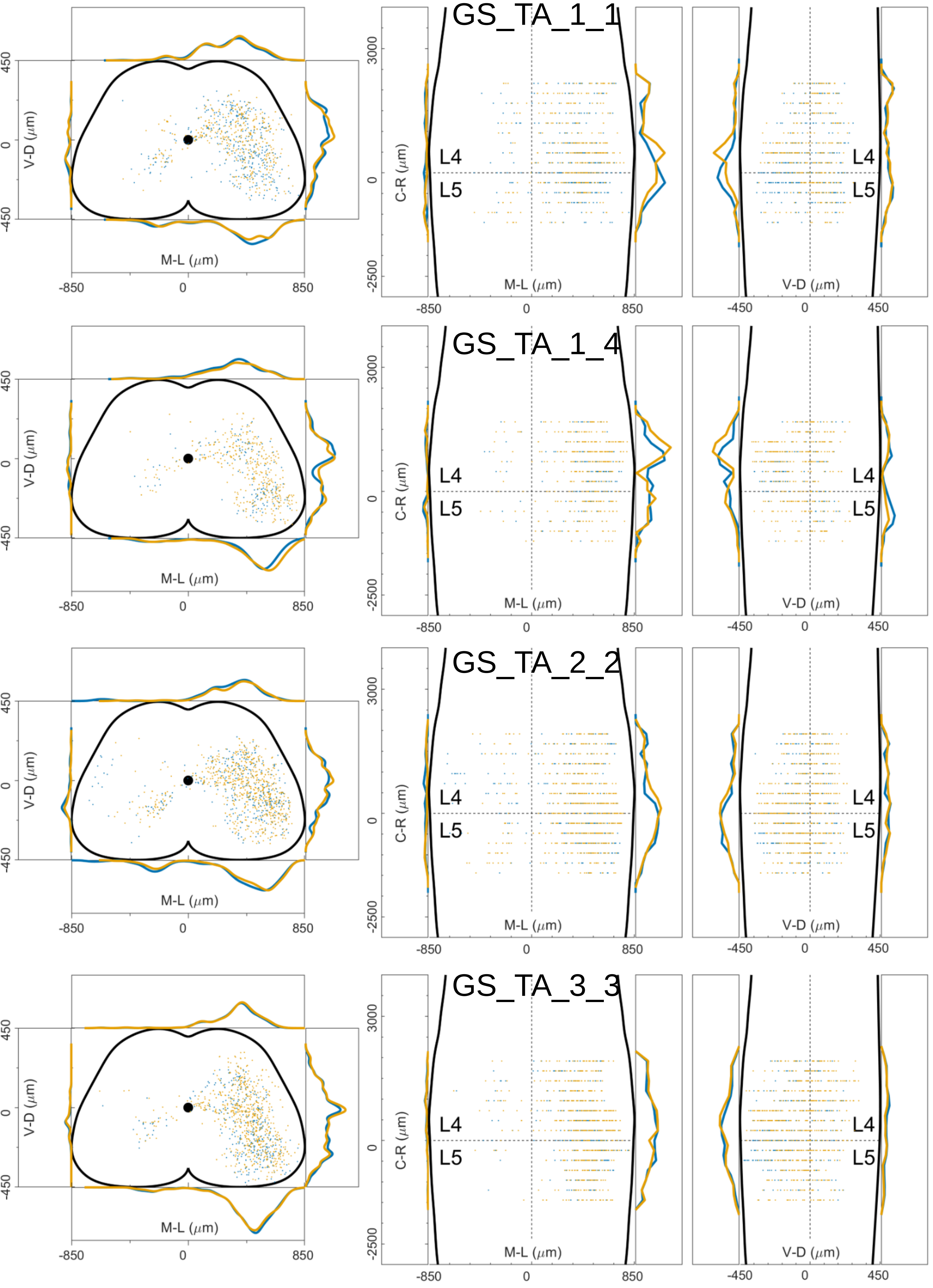

Figure 13 figure supplement 1

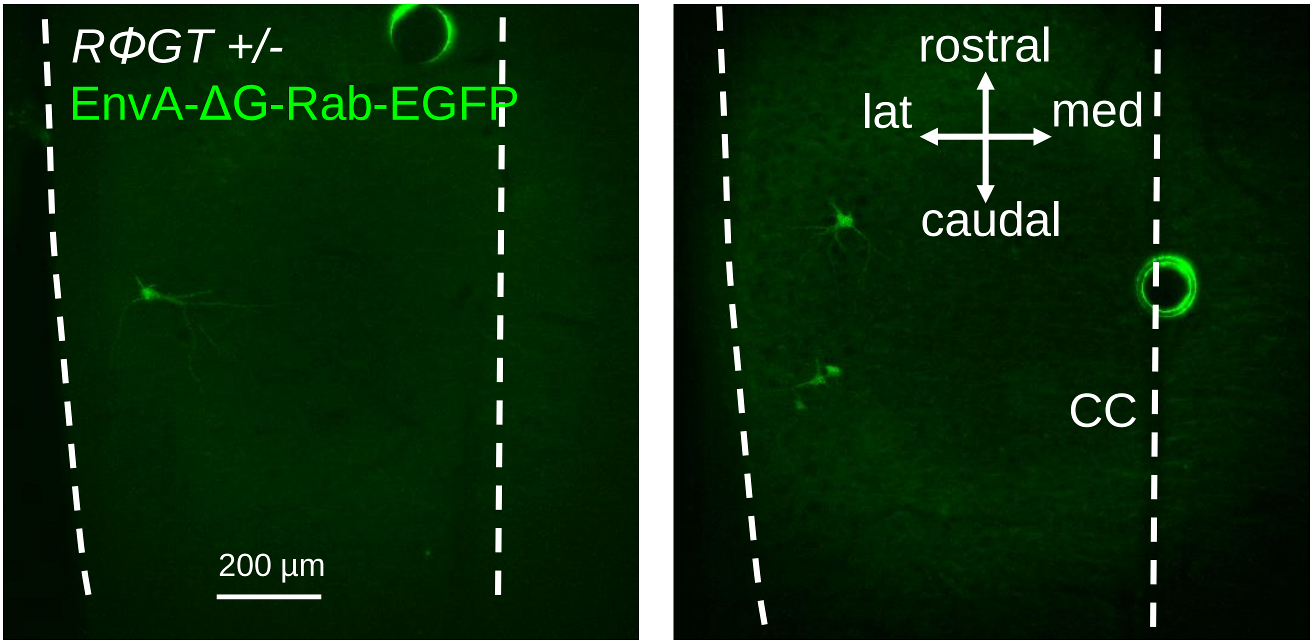

Figure 13 figure supplement 2

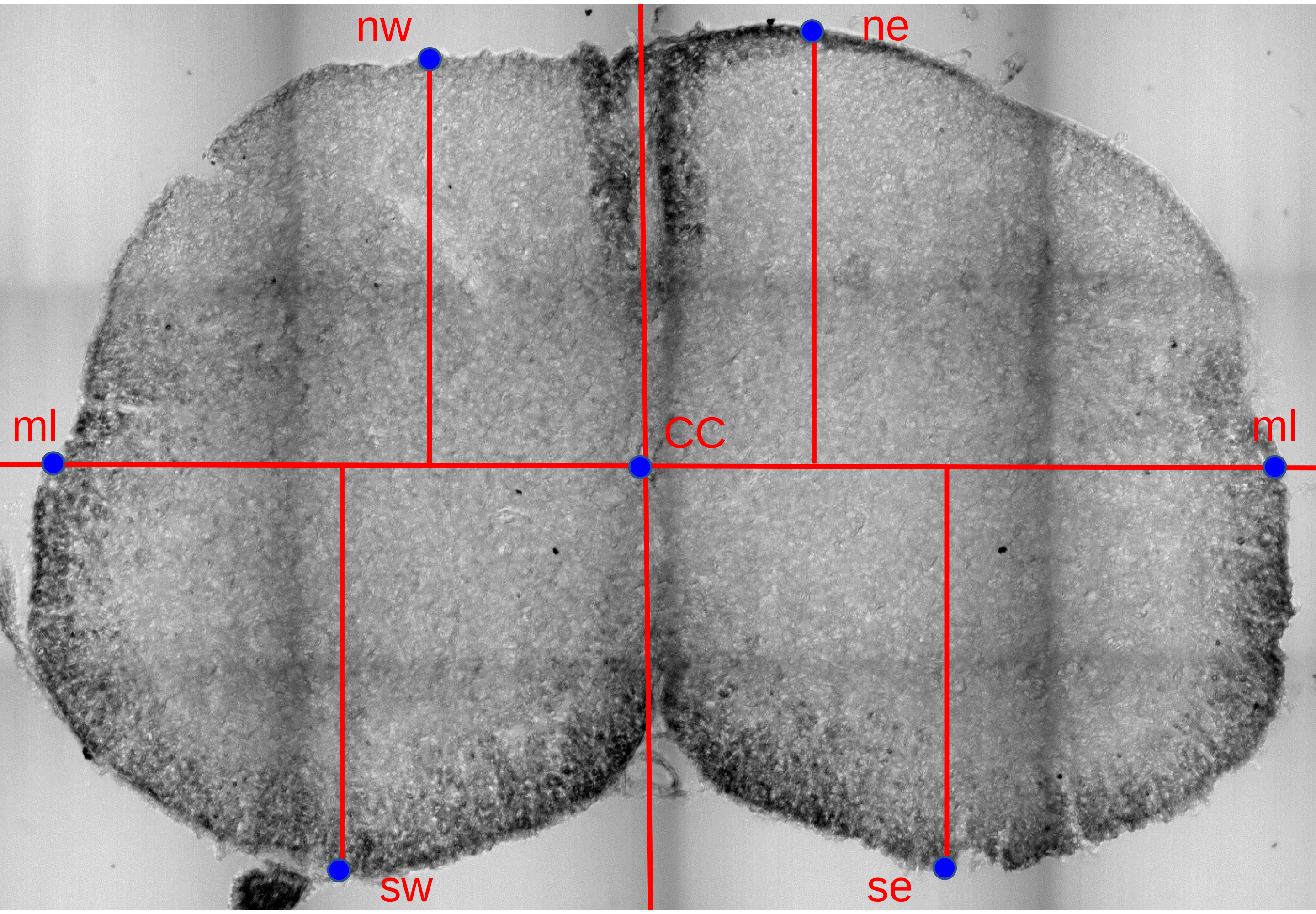
